# Expression of the CIC-DUX4 fusion oncoprotein mimics human CIC-rearranged sarcoma in genetically engineered mouse models

**DOI:** 10.1101/2023.09.26.559519

**Authors:** Peter G. Hendrickson, Kristianne M. Oristian, MaKenna R. Browne, Lixia Lou, Yan Ma, Dianna M. Cardona, Corinne M. Linardic, David G. Kirsch

## Abstract

CIC-DUX4 sarcoma (CDS) is a rare but highly aggressive undifferentiated small round cell sarcoma driven by a fusion between the tumor suppressor Capicua (CIC) and DUX4. Currently, there are no effective treatments and efforts to identify and translate better therapies are limited by the scarcity of tissues and patients. To address this limitation, we generated three genetically engineered mouse models of CDS (Ch7CDS, Ai9CDS, and TOPCDS). Remarkably, chimeric mice from all three conditional models developed spontaneous tumors and widespread metastasis in the absence of Cre-recombinase. The penetrance of spontaneous (Cre-independent) tumor formation was complete irrespective of bi-allelic CIC function and loxP site proximity. Characterization of primary and metastatic mouse tumors showed that they consistently expressed the CIC-DUX4 fusion protein as well as other downstream markers of the disease credentialing these models as CDS. In addition, tumor-derived cell lines were generated and ChIP-seq was preformed to map fusion-gene specific binding using an N-terminal HA epitope tag. These datasets, along with paired H3K27ac ChIP-seq maps, validate CIC-DUX4 as a neomorphic transcriptional activator. Moreover, they are consistent with a model where ETS family transcription factors are cooperative and redundant drivers of the core regulatory circuitry in CDS.

## Introduction

The increased use of molecular diagnostics in cancer care has improved our understanding of previously difficult to classify tumor types. Ewing-like sarcoma, or undifferentiated small round cell sarcoma, is an example of a tumor that is both rare and difficult to distinguish from other small round blue cell tumors^1^. Historically named for its histological and phenotypic similarities to Ewing sarcoma, Ewing-like sarcoma was a diagnosis of exclusion^2^. Recently, it was discovered that a majority of Ewing-like sarcomas harbor a reciprocal translocation at t(4;19)(q35;q13), leading to a CIC-DUX4 gene rearrangement^3^. The resultant fusion protein contains the DNA binding domain from CIC, a canonical tumor suppressor, and the transcriptional activation domain (TAD) of DUX4, an early embryonic pioneer factor^4^. The CIC-DUX4 oncoprotein behaves as a transcriptional activator, however, the mechanisms behind CIC-DUX4 mediated tumorigenesis remain unclear^5^.

Ectopic CIC-DUX4 expression transforms murine mesenchymal cells and osteochondrogenic progenitor cells with high efficiency^6^. Orthotopic allografts using these cells form aggressive undifferentiated small round cell tumors resembling human CIC-DUX4 driven sarcomas (CDS). To our knowledge, there is no existing autochthonous mammalian models of this malignancy. Prior work in our group has identified the importance of co-evolution of the tumor and tumor microenvironment when evaluating novel therapeutics and interventions such as radiotherapy^7^. In soft tissue sarcomas, radiation therapy is a mainstay of treatment, however, treatment efficacy varies across tumor types and histologies. Clinically, CDS is resistant to chemotherapy and radiation therapy^8,9^. Due in part to the extreme rarity of the disease and further confounded by the aggressive disease course, identification and clinical trials of new therapies are especially difficult. Therefore, a model system that recapitulates this tumor and enables insight into tumor initiation, progression, metastasis, immune evasion, and the response to treatment is of particular significance. In this work, we describe efforts to generate an autochthonous primary mouse model of CDS that mimics the human disease to serve as a preclinical platform for CDS therapy development.

## Methods

### Derivation of transgenic animals

All animal experiments were approved by Duke University Animal Care and Use Committee, protocol number A014-22-01. Ch7CDS mice were generated using homology-independent CRISPR/Cas9-mediated targeted integration. A donor vector was designed to insert loxP sites and sequence from the human DUX4 TAD at the c-terminus of the endogenous Cic locus on chromosome 7 (ENSMUG00000005442). Suitable sgRNAs were identified, validated in vitro, and then cloned into a CRISPR/Cas9 activator plasmid (Addgene #64073). G4 embryonic stem cells were co-transfected with the activator and donor plasmids and then selected in G418 media^10^. Embryonic stem (ES) cell clones containing the conditional knock-in were identified by PCR and validated with Sanger sequencing before injecting into donor ICR mouse morulae. Morulae were transplanted into female nurse ICR moms for gestation and delivery of chimeric pups.

Rosa26 Lox-STOP-Lox (LSL) Ai9-HA-CIC-DUX4 (R26-Ai9) mice were generated by subcloning an N-terminal 3x HA-tagged CIC-DUX4 fusion gene from Yoshimoto et al.^6^ into a Rosa26 targeting construct (Addgene #21714). The sequence verified construct was then transfected into ES cells and selected in G418 media. Clones containing the knock-in were identified by PCR, sequence validated, and then injected into donor ICR mouse morulae which were transplanted into female nurse ICR moms for gestation and delivery of chimeric pups.

Rosa26 LSL TOPO-HA-CIC-DUX4 (R26-TOPO) mice were generated by subcloning the LSL cassette from a Lox-STOP-Lox TOPO vector (Addgene #11584) into the Rosa26-Ai9-HA-CIC-DUX4 targeting construct. Transfection into ES cells, clonal selection, validation, and transplantation were carried out as described above.

### Genotyping

Genomic DNA (gDNA) was purified from ES cells and tail clips using the Quick-DNA Miniprep Kit (Zymo Research). Initial validation of target insertion was performed with primers designed to amplify across the 5’ and 3’ integration sites (Supplementary data 1-4). Several positive ES cell clones were then selected (indicated by red font) and verified by Sanger sequencing prior to expansion and morulae aggregation. To look for recombination, gDNA was also purified from tumors and tumor-derived cell lines as above and new primer pairs were designed to amplify across the entire region between the loxP sites (Supplementary data 4). PCR was performed using TaKaRa LA Taq and optimized for amplicon size.

### Derivation of tumor cell lines

Mouse soft tissue sarcoma cell lines were generated as described previously^11^. Tumors were resected using aseptic technique from humanely euthanized animals. Tumor tissue was enzymatically and mechanically digested by serial pipetting, washed, and resuspended in sterile PBS. The cell suspension was filtered through a 70 μm cell strainer, pelleted, and resuspended in DMEM containing 10%FBS and 1X GA-1000 antibiotic (standard growth media). Cells were plated at high density in tissue culture treated flasks and assessed for cell death the following morning. Viable adherent cells were maintained in standard growth media and passaged with 0.25% trypsin-EDTA.

### Western blot

Cells lines were maintained in standard growth media until 80% confluency. Using Trypsin-EDTA, the cells were lifted, collected in Hank’s Balanced Salt Solution, and then pelleted by centrifugation (300xg for 3 minutes). Lysates were made using RIPA buffer (supplemented with 1% SDS, HALT protease inhibitor (Thermo Fisher Scientific), and Benzonase) and quantified with Pierce BCA protein assay kits. Heat-denatured proteins were loaded onto a 10% Bis-Tris gel, run at 150-200v in 1x MES buffer, and then wet-transferred at 350mA for 1 hour at 4°C. Importantly, all above steps were completed same-day due to the transient and unstable nature of the fusion. The CIC-DUX4 fusion (∼260kD) was probed using an anti-HA antibody (Cell Signaling, 3724) at 1:1000 dilution and anti-DUX4 antibody (Abcam, ab124699) at 1:1000. Cre was detected using an anti-Cre antibody (Cell Signaling, 15036) at 1:1000 dilution with B-actin (Sigma, A1978) as a loading control. Images were acquired on a LI-COR Odyssey CLx and processed using the Image Studio Software.

### Immunohistochemistry

Tissue samples were fixed in 10% formalin/70% ethanol and embedded in standard paraffin blocks. Five micrometer sections were mounted and stained with hematoxylin and eosin (H&E) or antibodies. Chromogen amplification and detection of antibodies was performed using the Vectastain Elite ABC-HRP Kit and DAB Peroxidase Substrate Kit (Vector Labs). Antigen unmasking was performed with a citrate buffer pH 6.0 by a modified microwave retrieval method or boiling. Images were captured on a Leica inverted light microscope with DFC450 camera and processed using the Leica Application Suite. The following antibodies were used for immunohistochemistry: DUX4 (Thermo-Fisher, MA5-16147), HA (Cell Signaling, 3724), CD99 (Thermo-Fisher, bs-2523R), Pan-cytokeratin (abcam, ab9377), WT1 (Thermo-Fisher, PA5-116131), Sox10 (abcam, ab227680), Desmin (abcam, ab15200), CD34 (Thermo-Fisher, 14-0341-82). All slides were reviewed by an expert sarcoma pathologist (DC) at Duke University.

### RNA-sequencing

RNA was extracted and purified from flash frozen tumors/normal tissues and cells using a Qiagen RNeasy kit following the manufacturer’s instructions for fibrous tissue (Qiagen, Hilden, Germany). High quality RNA (RIN >7) was divided into triplicate from which 150bp paired-end, rRNA-depleted, libraries were made using the Illumina TruSeq RNA library Prep Kit (Illumina, CA, USA). Libraries were quantified using the KAPA Library Quantification kit (KAPA Biosystems, MA, USA), multiplexed, clustered onto flowcells, and then sequenced using an Illumina HiSeq 4000 sequencer (or equivalent platform) by GENEWIZ (Azenta, NJ, USA). Raw sequencing reads were trimmed using Trimmomatic v0.39 (ILLUMINACLIP:TruSeq3-PE-2.fa:2:30:10:2:keepBothReads LEADING:3 TRAILING:3 MINLEN:36) and then aligned to the mm10 reference genome using default parameters in STAR v2.7.10a. FeatureCounts (Subread v2.0.3) was used to compile a count table from sorted and indexed BAM files which was then loaded into DESeq2 for differential expression analysis. Raw data is available on GEO under the accession number GSE241369.

### ChIP-sequencing

ChIP-sequencing was performed in tumor-derived cell lines generated from Ai9CDS and TOPCDS mice using the Active Motif ChIP-IT High Sensitivity kit (ActiveMotif, CA, USA). Cells were seeded into 150mm dishes and cultured in standard growth medium. At 80% confluency, the cells were crosslinked in 37% formaldehyde (with methanol) for 15 minutes on plate and quenched with glycine. Washed cell pellets were manually lysed using a dounce homogenizer and the chromatin was fragmented using a Q125 sonicator (Qsonica, CT, USA) with the following settings: 25% amplitude, 30 seconds ‘ON’/30 seconds ‘OFF’ for 20 minutes total. Separate immunoprecipitation reactions using an Anti-HA tag antibody (Abcam; ab9110) or anti-Histone H3K27ac antibody (ActiveMotif, 39134) were setup for overnight immunoprecipitation reactions at 4°C. ChIP DNA was bound to Protein G agarose beads, column purified, and eluted. 150bp, paired-end, DNA libraries for sequencing were prepared using the TruSeq DNA library Prep Kit (Illumina, CA, USA) and quantified using the KAPA Library Quantification kit (KAPA Biosystems, MA, USA). Libraries were sequenced on an Illumina HiSeq 4000 sequencer (or equivalent platform) by GENEWIZ (Azenta, NJ, USA). Raw sequencing reads were trimmed using Trimmomatic v0.39 and then aligned to the mm10 reference genome using Bowtie 2 (-q -t --no-mixed --no-discordant). Duplicate reads were marked and removed using Picard tools v2.18.2 and peaks were called with MACS3 (-B -f BAMPE -g 1.87e9 -q 0.01). Peak files were filtered against the ENCODE blacklisted regions (https://github.com/Boyle-Lab/Blacklist/tree/master/lists) and then annotated using ChIPseeker v3.17^12^. De novo motif enrichment analysis on HA-CIC-DUX4 peaks was performed with HOMER v4.11^13^. Super Enhancers were called from H3K27ac peaks using ROSE and used as input for CRCmapper to map the core regulatory circuitry^14,15^. Raw data is available on GEO under the accession number GSE241370.

## Results

### Fusion of a human DUX4 C-terminal domain to endogenous Cic is sufficient to generate small round cell sarcomas in mice

To investigate the ability of CIC-DUX4 to generate tumors in vivo, we used CRISPR/Cas9 to insert the C-terminal domain (CTD) of human *DUX4* into exon 20-the most common breakpoint in human fusions-of the endogenous mouse *Cic* gene on chromosome 7^16^. This strategy was chosen over a pure endogenous fusion model due to the repetitive structure of Dux/DUX4 alleles and weak CTD conservation^17^. Here, prior to Cre exposure, mice express canonical Cic. After injecting the mice with Adenoviruses expressing Cre or by breeding with a tissue specific Cre driver, recombination between the loxP sites would excise the endogenous exon 20 and termination sequence creating and permitting the expression of a Cic-DUX4 fusion (Figure 1a). By this method, we sought to recapitulate the haploinsufficiency of *CIC* observed in naturally occurring CIC-DUX4 sarcomas. Screened and validated ES cell clones were successfully injected, and 38 viable chimeric pups were born to host ICR mother mice. As expected, chimeric animals showed a high contribution of donor genetic material (50-100%) and genotyping by tail clip at one week of age found that all 38 pups harbored an unrecombined transgenic allele (Supplementary data 1). Surprisingly, beginning at 3-weeks of age, the chimeric animals spontaneously developed large, and in some cases multifocal tumors involving the limbs, flank, back, abdomen, and head and neck region which rapidly grew and metastasized (Figure 1b). The animals were humanely euthanized when evidence of extensive tumor burden or illness was observed, and all discernable tumors were harvested for analysis. By 5 weeks of age, all 38 chimeric animals were dead (Figure 1c). Although none of the animals were exposed to Cre recombinase, PCR amplification across the loxP sites was indicative of tumor-specific recombination (Figure 1d). To further characterize, several tumors were evaluated using an immunohistochemical panel for small round cell sarcomas including: CD99, WT1, SOX10, Cyotkeratin (CK), CD34, and Desmin. Repeatedly, the tumors showed focal/patchy CD99 expression and strong WT1 expression consistent with human CDS (Figure 1e)^18–20^. Taken together, these data suggest that spontaneous (Cre-independent) CIC-DUX4 expression is sufficient to induce the formation of aggressive small round cell sarcomas in mice.

**Figure 1.**
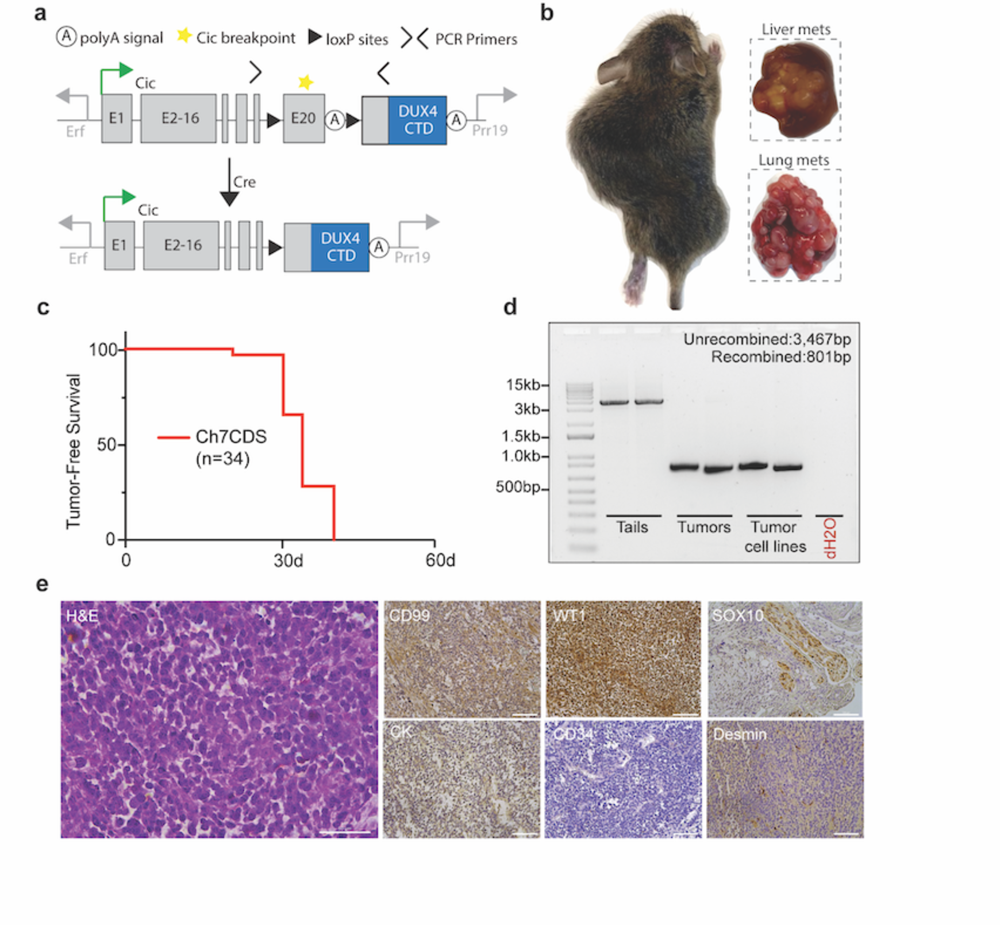
Fusion of a human DUX4 C-terminal domain to endogenous CIC is sufficient to generate small round cell sarcomas. a) Schematic of the Ch7CDS allele; Cre-LoxP recombination creates a *Cic*-DUX4 gene fusion at the endogenous *Cic* allele on chr 7. b) Gross images of primary and metastatic tumors from chimeric Ch7CDS mice. c) Kaplan-Meier survival curve of chimeric Ch7CDS animals. d) DNA gel of amplification products from PCR across the loxP sites in tails, tumors, and tumor-derived cell lines (n=2 each). In the tumor and tumor-derived cell lines, a ∼800bp PCR product is amplified consistent with loxP recombination. e) Representative images from an H&E stain (scale bar 50um) and immunohistochemistry (IHC) panel (scale bar 100um) on spontaneous tumors from Ch7CDS mice.

### CIC haploinsufficiency is not required for CIC-DUX4 sarcomagenesis

CIC is a highly conserved and canonical tumor suppressor that regulates MAPK effector gene expression^21^. In oligodendrogliomas, mutations in *CIC* are common and believed to be a key oncogenic event^22^. To test the necessity of CIC haploinsufficiency for CDS formation, we utilized a homologous recombination strategy to insert a 3x HA-tagged human CIC-DUX4 cDNA into the Rosa26 locus under control of a Lox-STOP-Lox cassette (Figure 2a). Here, the HA-CIC-DUX4 fusion is expressed without affecting the endogenous *Cic* alleles. In parallel with production of the Ch7CDS model, targeted ES cell clones were screened, validated, and implanted giving rise to 32 chimeric pups. Animals showed 50-100% chimeric contribution and genotyping at 1 week of age confirmed the presence of an unrecombined transgenic allele (Supplementary data 2). Again, beginning at 3 weeks of age, the chimeric Ai9CDS animals spontaneously developed tumors in the absence of Cre-recombinase and rapidly succumbed to disseminated disease. By 6 weeks of age, all animals had died naturally or were humanely euthanized (Figure 2b) with tumors histologically resembling CDS. PCR amplification across the loxP sites demonstrated recombination of the loxP sites in the tumors (Figure 2c). Based on the rapid tumor onset despite two intact *Cic* alleles, we conclude that CIC haploinsuffiency is not required for CIC-DUX4 sarcoma formation.

**Figure 2.**
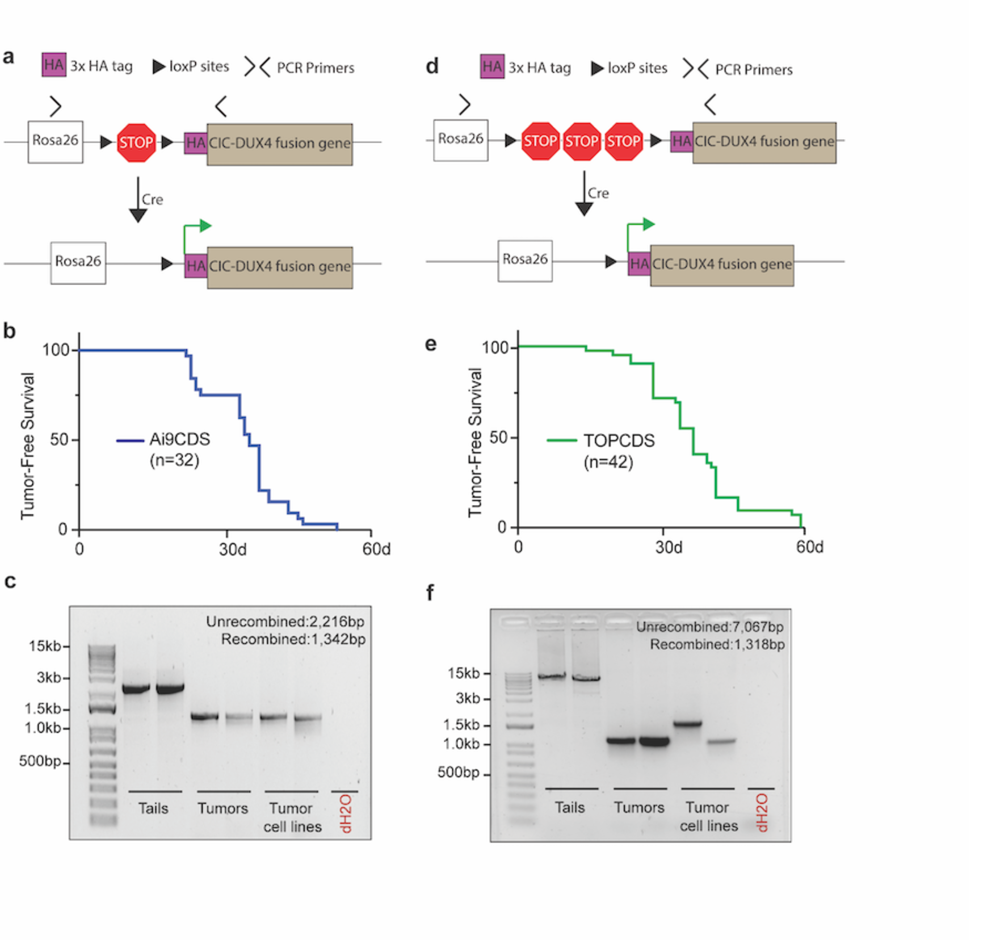
CIC haploinsufficiency is not required for CIC-DUX4 sarcomagenesis. a) Schematic of the Ai9CDS allele; Cre-LoxP recombination removes a stop cassette and permits expression of an HA-CIC-DUX4 fusion gene at the Rosa26 locus. b) Kaplan-Meier survival curve of chimeric Ai9CDS animals. c) DNA gel of amplification products from PCR across the loxP sites in tails, tumors, and tumor-derived cell lines (n=2 each). d) Schematic of the TOPCDS allele; Cre-LoxP recombination removes a stop cassette (containing multiple polyA sequences in tandem) and permits expression of an HA-CIC-DUX4 fusion gene at the Rosa26 locus. e) Kaplan-Meier survival curve of chimeric TOPCDS animals. f) DNA gel of amplification products from PCR across the loxP sites in tails, tumors, and tumor-derived cell lines (n=2 each).

### Extended STOP cassette does not prevent Cre-independent recombination

Genetic recombination is a fundamental biological process that is mediated, and exploited in genetic engineering, by the presence of direct repeat sequences such as loxP sites. In the Cre-Lox system, the likelihood and efficiency of recombination decreases with increasing distance between pairs of loxP sites^23^. To delay tumor onset, we engineered a third mouse model, TOPCDS, using the same strategy as the Ai9CDS mouse with one exception. In place of a short LSL cassette optimized for cloning, we utilized a long LSL cassette subcloned from a targeting construct used to control the expression of other potent oncogenes like Kras-G12D (Figure 2d)^24^. Compared to the ∼900bp LSL cassette in the Ai9 vector, the TOPO LSL cassette spans ∼5700 bp and contains two additional STOP sequences making it less susceptible to spontaneous recombination. 42 viable chimeric pups were born to host ICR mothers. Animals showed 50-100% chimeric contribution and genotyping at 1 week of age confirmed the presence of an unrecombined transgenic allele (Supplementary data 3). Beginning at 3 weeks, the animals once again developed spontaneous tumors in the absence of Cre which rapidly and widely metastasized. By 9 weeks of age, all 42 animals had died naturally or were humanely euthanized (Figure 2e) with tumors identical in appearance and histology to the first two models. Examination of loxP sites using PCR amplification showed tumor-specific recombination (Figure 2f). Thus, although the extended LSL cassette modestly increased tumor latency, it was not sufficient to prevent Cre-independent recombination.

### Tumors that arise in chimeric transgenic animals are driven by CIC-DUX4 in the absence of Cre recombinase

To validate that the tumors in our chimeric mice were driven by CIC-DUX4, in situ expression of the fusion protein was validated using immunohistochemistry. Here, antibodies against the N-terminal HA epitope tag as well as the C-terminus of human DUX4 were used to confirm expression of the fusion gene. Notably, in both Rosa26 models (Ai9CDS and TOPCDS), HA and DUX4 showed strong positive nuclear staining in the tumor cells but not in surrounding stroma or normal tissue (Figure 3a)^25^. As expected, Ch7CDS tumors exhibited strong nuclear DUX4 staining, and the HA epitope was not detected. Cre-recombinase was also not detected in any of the models. To rule out the possibility of transient expression from a cryptic Cre gene, PCR genotyping was performed on DNA isolated from ES cells, tails, and tumors from each of the three models. All assays for Cre, iCre, and strain-specific Cre were negative. To facilitate mechanistic and genomic studies, tumor-derived cells lines were generated from each of the three mouse models. Like the parent tumors, all three cell lines expressed the full-length CIC-DUX4 fusion oncoprotein in the absence of Cre-recombinase (Figure 3b). To confirm CIC-DUX4 transcriptional activity, RNA was harvested from tumors and tumor-derived cell lines for bulk RNA-sequencing. Using mouse KP (Kras^G12D^, p53^fl/fl^) sarcoma tumors^26^ and normal muscle as controls, all three CDS models strongly upregulated markers of CIC-rearranged sarcoma including ETV1/4/5, SHC3/4, DUSP4/6, and VGF further authenticating them as CIC-DUX4 sarcomas^5,20,27^ (Figure 3c).

**Figure 3.**
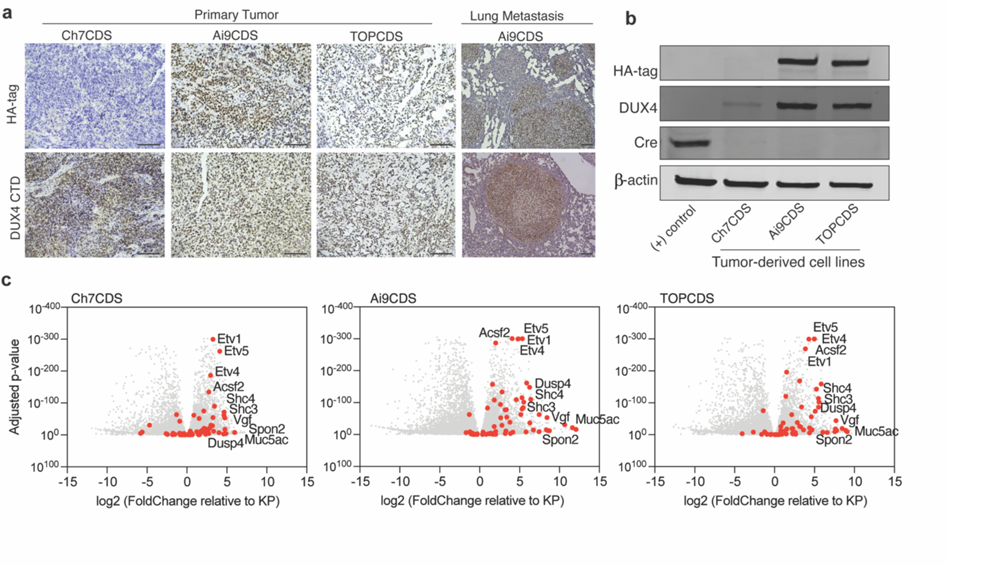
Mouse tumors express CIC-DUX4 and a transcriptional signature consistent with CDS. a) Representative images from HA-tag (top) and DUX4 (bottom) IHC in primary and metastatic tumors from Ch7CDS, Ai9CDS, and TOPCDS mice (scale bar 100um). b) Western blot on a Cre-expressing control cell line and tumor-derived cell lines probed for the HA-tag, DUX4 CTD, and Cre recombinase. DUX4 confirms the expression of a ∼260kD protein in all three CDS cell lines corresponding to the predicted size of the CIC-DUX4 fusion. Anti-HA is specific to the two epitope-tagged cell lines, however, all three CDS cell lines are negative for Cre. c) Volcano plots showing differentially expressed genes in CDS tumors compared to KP (Kras^G12D^, p53^fl/fl^) tumors. Red dots correspond to 65 genes highly and specifically expressed CIC-DUX4 target genes selected from human datasets^29,40^.

### CIC-DUX4 behaves as a neomorphic transcriptional activator

Several studies have inferred a neomorphic function for CIC-DUX4 as a direct transcriptional activator based on gene expression changes in transformed cell lines and ChIP-sequencing with non-specific antibodies^24,25^. Using the N-terminal 3x HA-epitope tag, CIC-DUX4 fusion gene specific ChIP-seq was performed in our Ai9CDS and TOPCDS cell lines revealing 4,861 and 5,057 high confidence binding sites, respectively (Figure 4a). Binding was most enriched at gene promoters (observed/expected:3-4, p<1e-70) and associated with genes involved in notable oncogenic signaling pathways including Hippo, Wnt, and PI3K-AKT (Figure 4b). Because fusion proteins have been known to acquire novel binding site capabilities, de novo motif enrichment analysis was performed on peaks shared between the two experiments (n=2,410). As anticipated, the most enriched motif matched the predicted binding site of CIC (CATT), however, a motif matching the consensus binding site for ETS transcription factors (GGAA) was also highly over-represented at HA-CIC-DUX4 binding sites (Figure 4c). To understand the effect of CIC-DUX4 binding on local chromatin, ChIP-seq for H3K27ac was performed in parallel. Predictably, HA-CIC-DUX4 peaks strongly co-localized with H3K27ac in keeping with its pre-conceived behavior as a transcriptional activator (Figure 4d). Lastly, we sought to identify other core transcription factors that may co-operatively regulate the transcriptional circuity in CDS. To do so, Ai9CDS and TOPCDS cell line H3K27ac ChIP-seq datasets were used to define Super Enhancers (SE) which were then interrogated for known transcription factor motifs. This analysis identified 11 transcription factors common between both cell lines all of which, with the exception of Tead1 and Creb3l2, are highly upregulated in mouse and human CIC-DUX4 sarcomas (Figure 4e)^29^. Remarkably, 4 of the 11 predicted genes are ETS transcription factors (i.e. Etv5, Etv4, Etv1, and Ets1) adding to a body of literature which has previously implicated these factors as critical drivers of CIC-DUX4 sarcomagenesis^5,28,30^.

**Figure 4.**
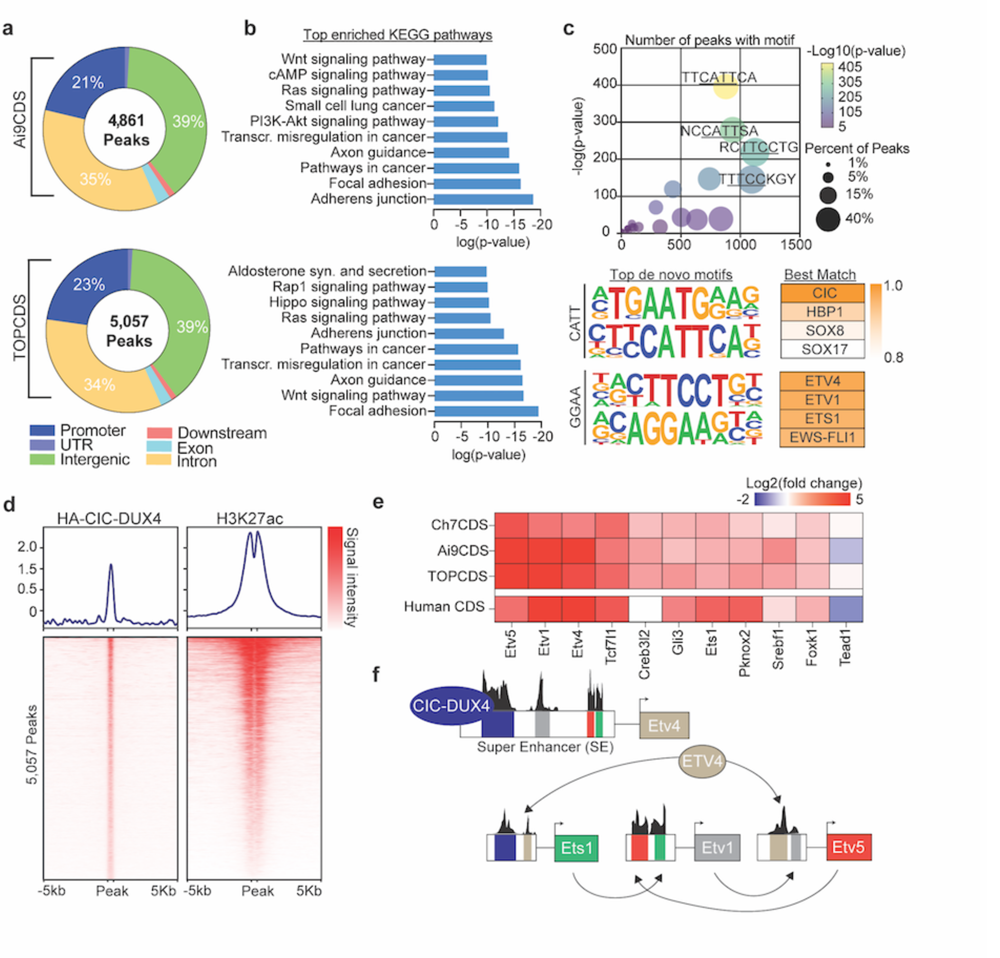
CIC-DUX4 behaves as a neomorphic transcriptional activator. a) Pie charts corresponding to the genomic location of all HA-CIC-DUX4 peaks in tumor cell lines from Ai9CDS and TOPCDS mice. b) Top enriched KEGG pathways based on all genes associated with a HA-CIC-DUX4 peak. c) Dot plot (top) showing the top 25 most enriched de novo motifs identified in 2,410 shared HA-CIC-DUX4 peaks. The top four de novo motifs match the predicted binding sites for CIC (AATG/CATT) and ETS-family transcription factors (TTCC/GGAA). d) Ranked heatmaps showing relative read coverage from HA-CIC-DUX4 and H3K27ac ChIP across all 5,057 HA-CIC-DUX4 peaks in TOPCDS cells. e) Heatmap displaying the log2 fold change (tumor/KP) for all 11 transcription factors predicted to be ‘core regulators’ of the CDS transcriptional circuitry. On the bottom row, log2 fold change in human CDS relative to ES^29^ is included. f) Summary of the results from ‘CRCmapper’ highlighting the interconnected auto-regulatory loop between CIC-DUX4 and ETS transcription factors.

## Discussion

CIC-DUX4 sarcoma (CDS) is a rare and highly aggressive undifferentiated small round cell sarcoma affecting adolescents and young adults^31^. Despite its clinical and morphological similarities to Ewing sarcoma, CDS is insensitive to conventional Ewing Sarcoma therapies translating to high rates of relapse and low survival^32^. To improve patient outcomes, the mechanisms underlying tumor initiation, maintenance, and metastasis need to be understood. One obstacle to acquiring this knowledge is the scarcity of primary tissues and patients necessary for developing and testing specific hypotheses. To overcome this, an animal model with an intact immune system that emulates the aggressive properties would be a valuable resource to the field. To this end, we worked to develop a genetically engineered mouse model of CDS. Surprisingly, the chimeric animals from all three models, irrespective of *Cic* haploinsufficiency, developed aggressive tumors and metastatic disease that was uniformly lethal between 3-9 weeks of life. All tumors were nearly identical to human CDS in appearance and expressed a CIC-DUX4 fusion gene in the absence of Cre-recombinase. Although spontaneous (Cre-independent) recombination has been reported in other genetically engineered mouse models with conditional oncogenes, the event rate was low and tumors were specific to a few susceptible tissues and organs^33^. In our models, the penetrance was complete and widespread underscoring a strong positive selective pressure for the CIC-DUX4 fusion oncogene. Consistent with prior work, our RNA-seq and ChIP-seq data suggests that fusion of the DUX4 CTD converts the CIC transcriptional silencer into a potent transcriptional activator. Likely, this is mediated by the recruitment of P300/CBP and/or other histone acetyltransferases to genes typically silenced by CIC^34^. The genes most responsible and essential for the aggressive properties of CIC-DUX4 tumors remains to be determined. Our analysis of the transcriptional network activated in mouse CDS points to ETV1, ETV4, ETV5, and ETS1 as conserved targets of CIC-DUX4 and, more importantly, as key cooperative (and potentially redundant) regulators of the CDS transcriptional circuitry (Figure 4f). The role of ETV4 in CDS has been investigated before but the results are inconsistent. For example, in a transgenic zebrafish model of CDS, genetic loss of ETV4 impairs tumor formation^30^. Conversely, ETV4 knockdown in transformed CIC-DUX4-expressing NIH 3T3 cells had no effect on cell viability and tumor growth but did impair metastatic potential^28^. One explanation for the discrepancy may be a divergence in PEA3 subfamily redundancy during evolution from zebrafish to human^35,36^. Another possibility relates to the species-specific distribution of ETS binding sites which has forestalled numerous attempts to model Ewing Sarcoma in mice^37^. To dissect the independent and overlapping roles of ETS1 and PEA3 subfamily genes in CDS, future work will use CRISPR/Cas9 to systematically and combinatorially delete these genes in cell lines and tumors. Of further interest is whether forced overexpression or stabilization of these same factors could be toxic to CDS cells in keeping with the ‘Goldilocks phenomenon’ described in Ewing Sarcoma^38,39^. Lastly, these results indicate that CDS, unlike Ewing Sarcoma, can be modeled in the mouse which is an important breakthrough for the sarcoma research community. However, innovative solutions to prevent or overcome the effects of spontaneous recombination will be required to generate a model that is both sustainable as well as spatially and temporally controllable for experimentation and discovery.

## Acknowledgements

This work was supported by grants from the Department of Defense (W81XWH-22-1-0454) to DGK, Alex’s Lemonade Stand Foundation to CML, the National Cancer Institute (7R35CA197616) to DGK and (1R38CA245204) PGH, and the American Society of Radiation Oncology (852948) to PGH.

---

**Supplementary data 1.**
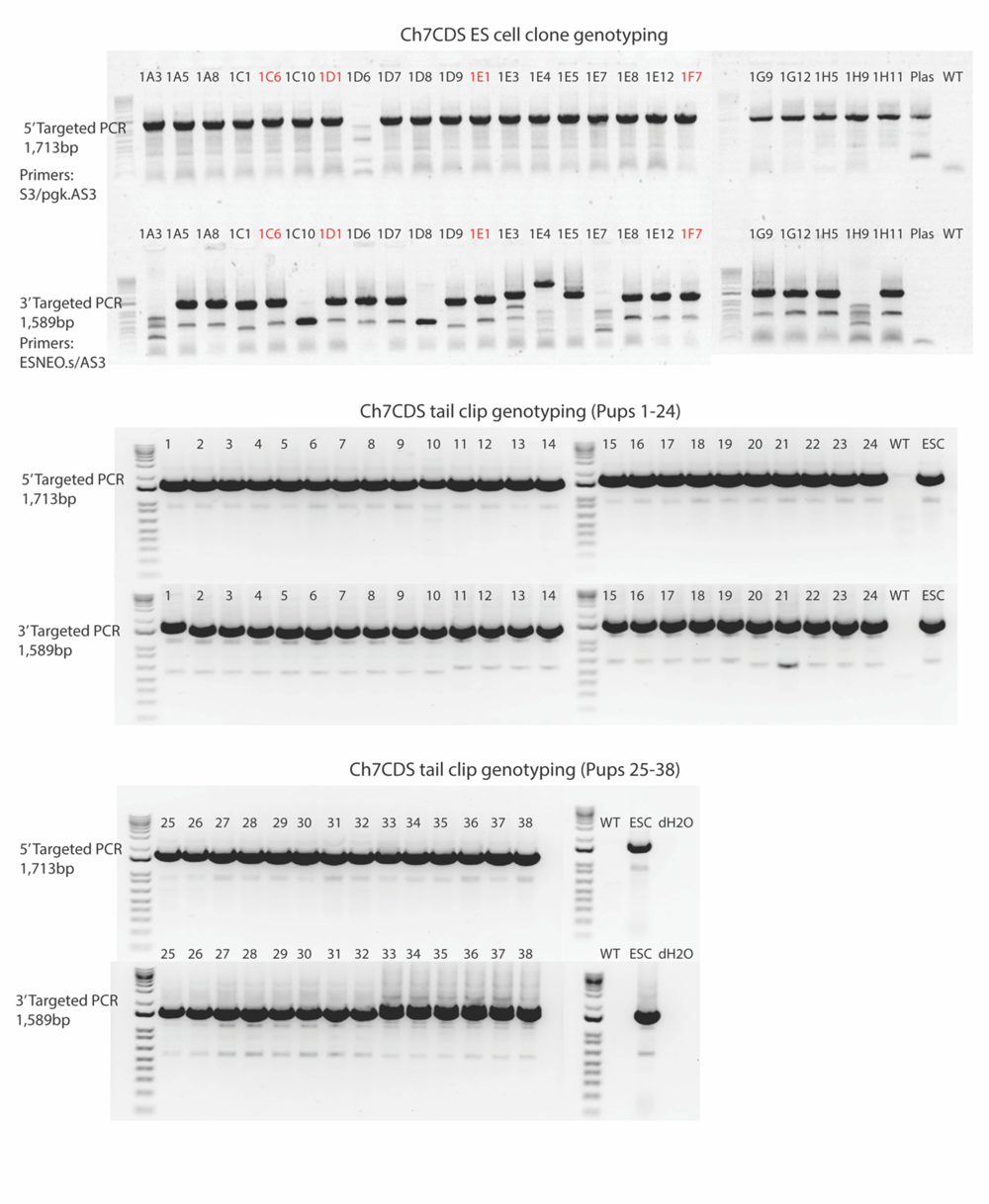

**Supplementary data 2.**
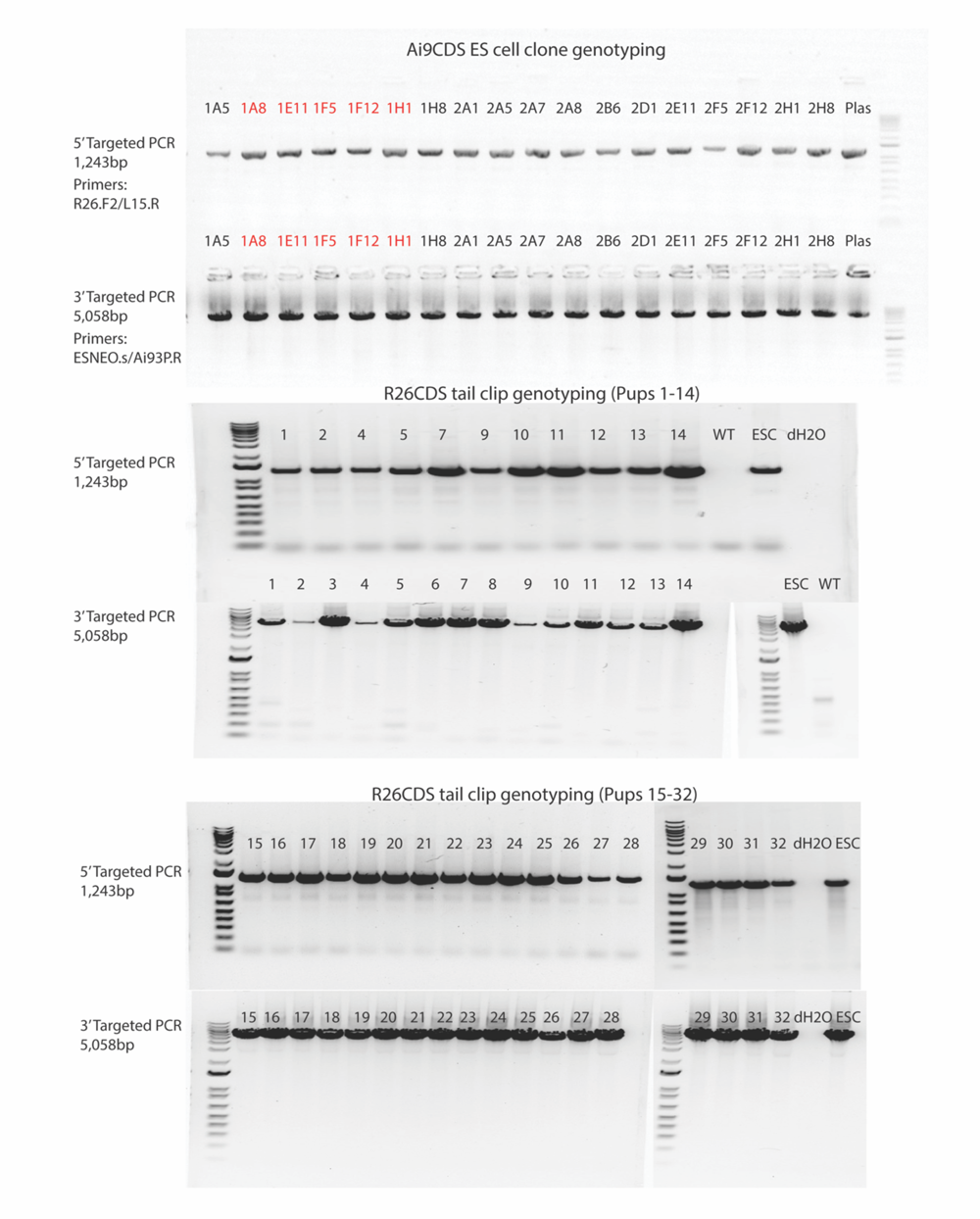

**Supplementary data 3.**
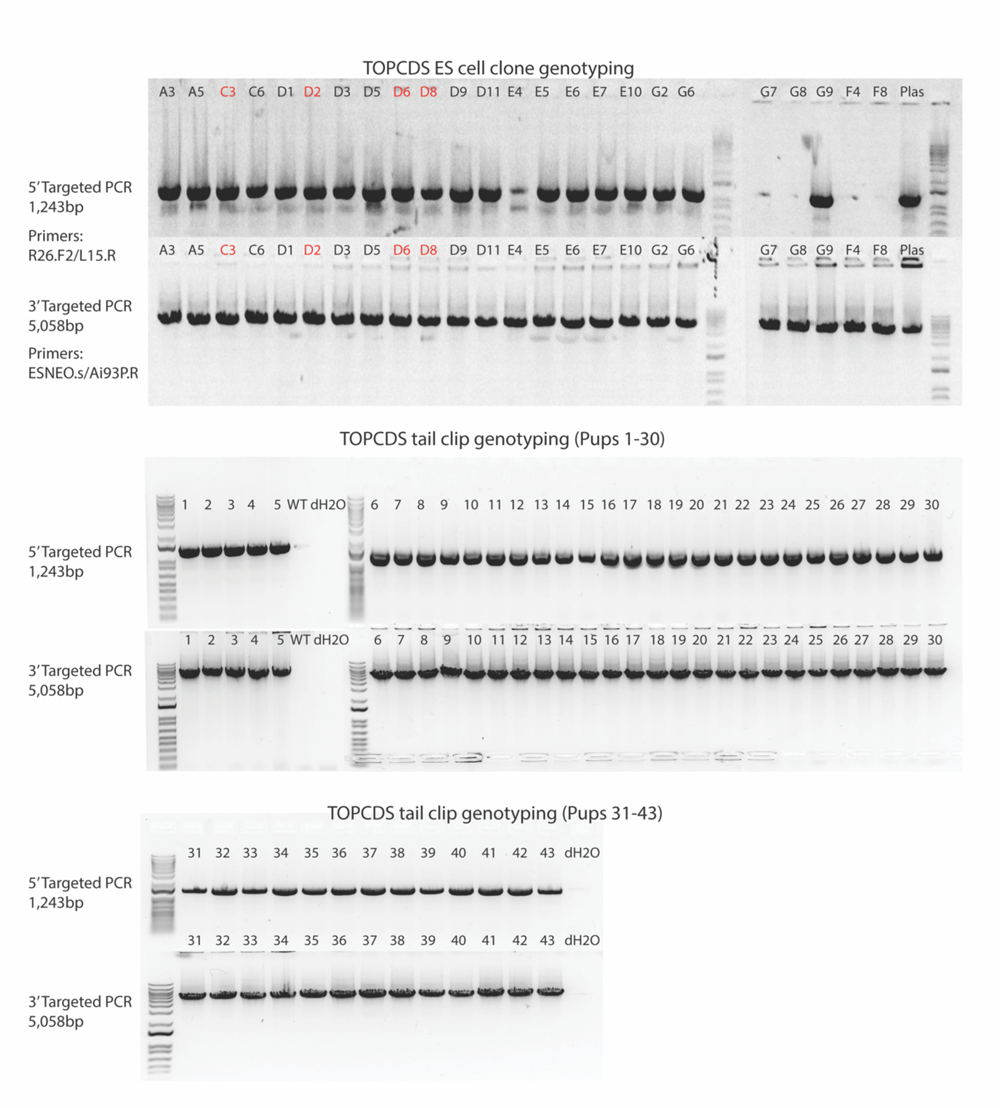

**Supplementary data 4.**
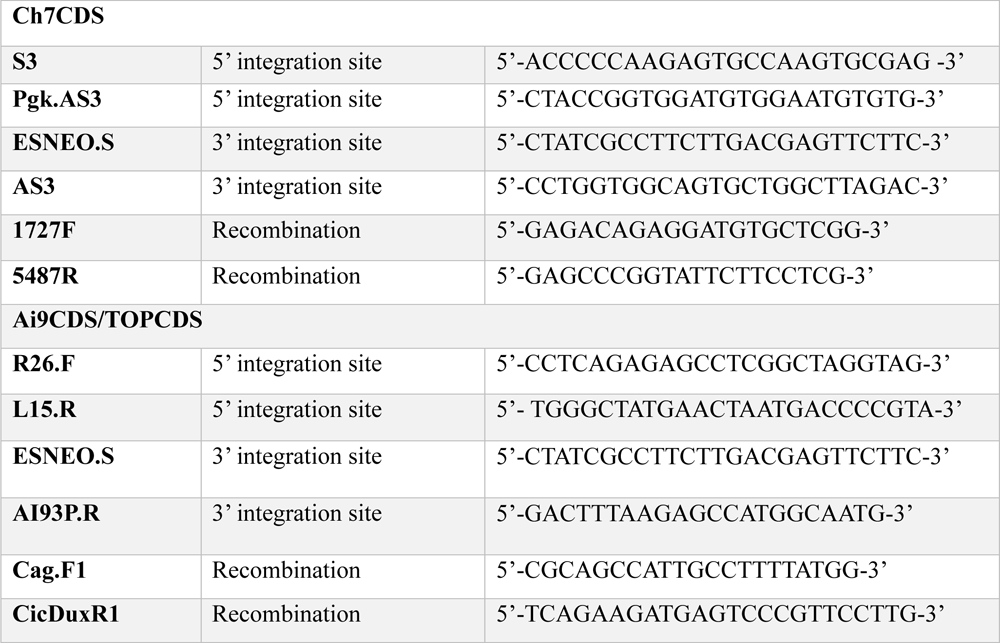

